# Sample size considerations for species co-occurrence models

**DOI:** 10.1101/2024.09.20.614180

**Authors:** Amber Cowans, Albert Bonet Bigatà, Chris Sutherland

**Author notes:** Joint lead authors (Amber Cowans, Albert Bonet Bigatà). Open research statement: Readers can access underlying simulated data and code at https://osf.io/puvck/. All data/code will be permanently archived on this Open Science Framework repository.

## Abstract

Multispecies occupancy models are widely applied to infer interactions in the occurrence of different species, but convergence and estimation issues under realistic sample sizes are common. We conducted a simulation study to evaluate the ability of a recently developed model to recover co-occurrence estimates under varying sample size and interaction scenarios while increasing model complexity in two dimensions: the number of interacting species and the number of covariates. Using both standard and penalized likelihood, we demonstrate that the ability to quantify interactions in species occupancy using this model is highly sensitive to sample size, detection probability and interaction strength. In the simplest scenario, there is high bias in the interaction parameter (used for co-occurrence inference) with less than 100 sites at high detection, and 400-1000 sites at low detection, depending on interaction strength. Strong co-occurrence is detected consistently above 200 sites with high detection probabilities, but weak co-occurrence is never consistently detected even with 2980 sites. We demonstrate that mean predictive ability of the co-occurrence model is less affected by sample size, with low bias in derived probabilities at 50 sites. Our results highlight that while occupancy patterns are often robust to sample size limitations, reliable inference about co-occurrence demands substantially larger datasets than many studies currently achieve. We caution the interpretation of model output in small datasets or when co-occurrence is expected to be weak, but show methods are suitable to quantify strong co-occurrence in larger datasets and generate predictions of site occupancy states.

## Introduction

Quantifying and predicting species-habitat and species-species associations is fundamental to ecology, where such processes underpin spatial distributions and biodiversity (Linnell and Strand 2000). When studying factors affecting species distributions, it is common practice to divide study areas into distinct sites and record species presence or absence (i.e., site occupancy: MacKenzie et al. 2002; Kéry and Royle 2015). These data are often collected using standardized survey methods such as camera trapping, acoustic monitoring, or direct field observations, leading to repeated detection of multiple species across space and time (Kerry et al. 2022). Occupancy models applied to such detection-non-detection data are now routinely used to study spatial distributions, especially in low-density populations (MacKenzie et al. 2002, 2005; Burton et al. 2015). In its simplest form, the classical occupancy model accounts for imperfect detection of the true but partially observable occupancy state by combining two distinct probability models: (1) a state model for occupancy, the underlying ecological variable of interest, and (2) an observation model linking imperfect detection to the latent state (MacKenzie et al. 2002). Although associations with habitat features are fundamental to understanding spatial distributions, accounting for non-independence in the occupancy of two or more species, which may arise from avoidance or attraction of one species towards another, is increasingly of interest (Burton et al. 2015; Rota et al. 2016).

Quantifying species co-occurrence using occupancy models is challenging, requiring either strong assumptions of interaction direction (Waddle et al. 2010) or prohibitively large data sets (MacKenzie et al. 2004). Recently, however, a multispecies occupancy model for two or more interacting species was proposed (Rota et al. 2016, ‘co-occurrence’ model from here). This framework offers the ability to explicitly quantify whether the presence of one species affects the occupancy (or detection) of another, over and above the effect of covariates included in the species-specific model (Rota et al. 2016; Kéry and Royle 2020) and perhaps most interestingly, whether these associations vary over space (Rota et al. 2016; Twining et al. 2022). Although one of several methods available for inferring species co-occurrence, the ease of implementation in the R package **unmarked** (Kellner et al. 2023), and the unique ability to model interaction strengths as a function of covariates, makes this model a very popular choice (cited 293 times as of April 2025).

The co-occurrence model was developed and motivated using empirical data from 1906 camera trap sites and 42,556 trap days (Rota et al. 2016) - a level of effort that the majority of studies applying the model fail to replicate. For example, we reviewed the literature citing Rota et al. (2016) and found that 59% of the studies that apply the model had fewer than 200 sites, and 35% had fewer than 100 sites (Appendix S1: Figure S1). Given that even standard single-species occupancy models show significant bias and low precision at 100-200 sites (Guillera-Arroita et al. 2010), these sample sizes are concerning, as extensions that increase model complexity are likely subject to similar sample size-related performance reductions. Moreover, convergence and estimation issues have been highlighted for the co-occurrence model in smaller data sets, i.e., between 50 and 400 sites (Clipp et al. 2021; Lauret et al. 2023). In these cases, a penalized likelihood approach has been developed which constrains unreasonably large parameter estimates and allows more complex models to be applied in these scenarios (Clipp et al. 2021). However, the trade-off for the variance reduction is estimator bias and unreliable standard errors, which may hinder inference on the existence and strength of species co-occurrence (Clipp et al. 2021). Despite several suggestions that estimation issues exist (Kéry and Royle 2020) and the increasing use of these models to inform ecological understanding and conservation management (Blanchet et al. 2020), the effects of sample size and model complexity on general model performance have not been investigated in sufficient detail. Given the rapid uptake of the co-occurrence model and a corresponding widespread tendency to interpret statistical associations as ecological interactions (Blanchet et al. 2020), this is an important and potentially problematic knowledge gap.

Here, using an extensive simulation study, we evaluate the co-occurrence model’s ability to recover known co-occurrence patterns under different sample sizes (*N* = 20 to *N* = 2980 sites) and co-occurrence scenarios. We first test sample size-based performance under the simplest parameterization of the model, i.e., a null model with two interacting species. We then increase model complexity in two dimensions: the number of interacting species and the number of covariates on the species interaction term. For each scenario, we implement both the standard and penalized likelihood approaches to directly compare the inference-prediction trade-off.

## Methods

### Co-occurrence model description

We first provide an overview of the multispecies occupancy model for two or more interacting species (Rota et al. 2016, i.e., the ‘co-occurrence’ model). In the standard occupancy model, across *j* = 1, …, *J* sampling occasions at *i* = 1, …, *N* sites, the detection (*y*_*i j*_ = 1) or non-detection (*y*_*i j*_ = 0) of a species is modelled as a Bernoulli random variable, conditional on the underlying latent occupancy state *z*_*i*_, where *z*_*i*_ = 1 if the species is present and *z*_*i*_ = 0 if it is absent (MacKenzie et al. 2002).

In the multispecies extension, each site is occupied or unoccupied by each of *s* = 1, …, *S* species and can therefore take on one of 2^*S*^ possible states. Species detection is modelled as a Bernoulli random variable conditional on the occupancy state of species *s* at the site *i* (*z*_*si*_) and species-specific detection probability *p*_*si j*_:

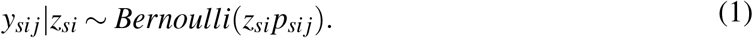

The site-specific occupancy state, **z**_*i*_, is an *S*-dimensional vector describing the true occupancy state of each species, **z**_*i*_ = {*z*_1*i*_, *z*_2*i*_, *z*_3*i*_, …, *z*_*Si*_}), and is modelled as a multivariate Bernoulli random variable with the expected probability vector *ψ* (Rota et al. 2016; Kéry and Royle 2020). With two species, A and B, a site has 4 possible states: both are absent [00]; A is present and B is absent [10]; A is absent and B is present [01]; and both are present [11]. The corresponding state vector is ***z***_*i*_ = {*z*_00,*i*_, *z*_10,*i*_, *z*_01,*i*_, *z*_11,*i*_}, and the corresponding probability vector is *ψ*_*i*_ = {*ψ*_00,*i*_, *ψ*_10,*i*_, *ψ*_01,*i*_, *ψ*_11,*i*_}, respectively (Rota et al. 2016). In the two-species case, the probability mass function of the multivariate Bernoulli distribution can be represented as a set of three ‘natural parameters’ which can be modelled as a function of parameter-specific covariates **X**:

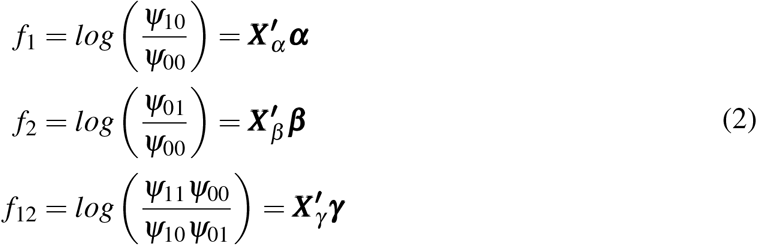

where *α, β, γ* are the coefficients for the corresponding covariates ***X*** ^***′***^.

The natural parameters in Equation 2 have a biological interpretation in terms of the log odds of occupancy: *f*_1_ and *f*_2_ are termed ‘first-order’ natural parameters and represent the log odds of species occurrence conditional on the absence of all other species included in the model; *f*_12_ is termed the ‘second order’ natural parameter and is the change in the log odds of species occurrence when the other interacting species is present, relative to when it is absent (Rota et al. 2016). There is, therefore, total pairwise independence between the occupancy of two species when *f*_12_ = 0, whereas *f*_12_*≠* 0 indicates positive or negative associations, depending on the sign, in species occurrence over and above the effects of covariates included in the first-order terms. In total, models can have up to 2^*S*^ *−* 1 natural parameters. However, in practice, it is common to only model pairwise interactions and set any natural parameter above second-order to zero (Kéry and Royle 2020). These natural parameters can be modelled as a function of covariates for inference about predictors of species occurrence (*f*_1_, *f*_2_) and co-occurrence (*f*_12_). From the natural parameters, the expected state-specific probabilities, e.g., *ψ*_00_, *ψ*_10_, *ψ*_01_, and *ψ*_11_, can be recovered using the multinomial logit link function (Rota et al. 2016). These probabilities are referred to as the ‘general parameters’. Predictions of both the marginal occupancy probabilities (i.e., the probability of species presence irrespective of occupancy of the others) and conditional occupancy probabilities (i.e., the probability of species presence conditional on the presence or absence of the others) can be derived from the general parameters using basic probability rules (Rota et al. 2016).

### Data simulation

We evaluated the performance of the co-occurrence model by fitting the model to simulated datasets and evaluating its ability to recover known data generating values, considering 18 scenarios of increasing complexity (Table 1). For each scenario, we simulated and analyzed 1000 co-occurrence datasets with three *J* = 3 sampling occasions (*visits* hereafter) and 11 distinct sample sizes, i.e., number of sites *N*, increasing from 20 to 2980 (*exp*(3) → *exp*(8) in increments of *exp*(0.5)). This range of sample sizes was chosen to reflect published studies from data-scarce, *N* = 20, to data-rich, *N* ≈ 3000 (Appendix S1: Figure S1).

**Table 1:** Features of the 18 simulation scenarios considered. Each simulation scenario was tested at 11 unique sample sizes (from 20 to 2980).

| Features varied | Number of scenarios |  |  |
| --- | --- | --- | --- |
|  | Null models for two species | Network complexity | Covariate complexity |
| **Interaction strength** | 3 (weak, moderate, strong) | Constant pairwise (moderate, moderate-strong, strong) | Constant pairwise (moderate, moderate-strong, strong) |
| **Interaction direction** | 2 (positive, negative) | Constant pairwise | Constant pairwise |
| **Detection probability** | 2 (high, low) | 1 (high) | 1 (high) |
| **Covariates** | 1 (none) | 1 (none) | 3 (1 unique, 1 shared, 1 unique + 1 shared) |
| **Species** | 1 (2 species) | 3 (3, 4, 5 species) | 1 (2 species) |
| **TOTAL** | 12 scenarios | 3 scenarios | 3 scenarios |

#### Null models for two interacting species: effect of sample size and detection probability

We first explored the effect of sample size using the simplest model structure with two interacting species (*S* = 2) and no covariates (null model) under high (*p*_1*i j*_ = *p*_2*i j*_ = 0.5) and low (*p*_1*i j*_ = *p*_2*i j*_ = 0.25) per-visit species-specific detection probabilities (Equation 1). These were kept constant across all site-visit combinations, so that the probability of detecting each species at least once across all three visits was equal to 0.88 and 0.58 for high and low detection scenarios, respectively. For each detection scenario, we considered six combinations of interaction strength (strong, moderate, weak) and direction (negative, positive) resulting in 12 scenarios. In Equation 2, we define a strong interaction as *γ*_0_ = *±*1, moderate as *γ*_0_ = *±*0.6, and weak as *γ*_0_ = *±*0.2.

#### Null models for more than two species: effect of network complexity

We then considered a further three scenarios increasing the number of species from *S* = 2 to *S* = 3, 4, 5 (network complexity), assuming constant interactions and no covariate effects. When adding more species to the network, we limited the maximum order of interactions to two (i.e. pairwise) as is common in the literature (Kéry and Royle 2020) and kept the interaction values between a species pair constant across all scenarios irrespective of *S* (e.g. *f*_12_ = 1 when *S* = 3, 4, 5). We set the pairwise interaction terms to moderate (0.6), moderate-to-strong (0.8), and strong (1) values for positive and negative second-order natural parameters. This allowed us to minimize the confounding effects of weak interactions that are known to be difficult to detect, and instead focus on evaluating model performance under more favorable conditions. By ensuring interactions were strong enough to be detectable, we could more clearly observe how additional challenges (i.e., increased complexity) impacted power, bias, and convergence. Values were chosen across the range of interaction directions and strengths whilst ensuring simulated data did not result in observations of all zeros or ones for any species (i.e. complete separation). All data-generating values can be found in (Appendix S1: Table S1). In addition, we fixed a high per-visit detection probability (*p*_*Si j*_ = 0.5) across all network scenarios to ensure that variation in performance was not driven by low detectability.

#### Covariate models for two species: effect of covariate complexity

Finally, we considered three additional scenarios where we altered the covariate structure for the second-order natural parameters in a model with only two interacting species (Equation 3). In all of these scenarios, we simulated the first-order natural parameters as a function of one covariate (*X*_*A*_ for *f*_1_, *X*_*B*_ for *f*_2_). In the first scenario, we simulated the interaction parameter (*f*_12_) as a function of one unique covariate (*X*_*C*_). In the second scenario, we simulated the interaction parameter as a function of one shared covariate (*X*_*A*_). In the third scenario, we simulated the interaction parameter as a function of two covariates, one unique and one shared (*X*_*C*_ and *X*_*A*_ respectively):

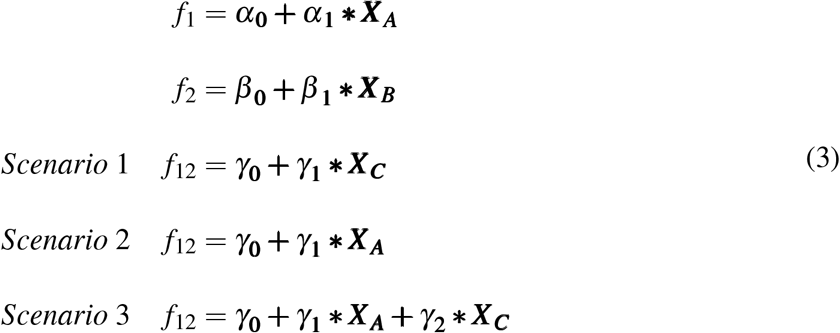

All covariates were continuous random variables generated from a standard normal distribution. As with the network scenarios, we constrained per-visit detection probabilities to *p*_*Si j*_ = 0.5 and interaction strengths to 0.6, 0.8 and 1 (Appendix S1: Table S1).

In all the 18 scenarios described above, we adjusted the values for the intercepts in the first-order natural parameters to ensure consistency in marginal species occupancy across and within scenarios (*ψ*_*S*_ ≈ [0.4 *−* 0.6]). For example, for the two-species null model scenarios, we set *α*_0_ = *β*_0_ = *−*0.4 when the interaction was positive and *α*_0_ = *β*_0_ = 0.1 when the interaction was negative, irrespective of interaction strength. These adjustments avoided patterns of systematic biases from either too many or too few instances of species occurrence or co-occurrence affecting parameter identifiability. We also discarded simulated datasets that had no observations for any of the species.

### Model performance

We fit the co-occurrence model to our simulated data using both likelihood (LL) and penalized likelihood (PL) methods in **R** (R Core Team 2023) using the `occuMulti()` and `optimizePenalty()` functions for LL and PL, respectively, from the **unmarked** package (Kellner et al. 2023). For PL methods, we estimated the optimal penalty value 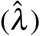 using the default settings of the fitting function, except when the standard likelihood approach failed to estimate the Hessian matrix. In these cases, we selected the optimal penalty value using cross-validation over the range of penalty values used in Clipp et al. (2021). When 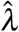 was at either the maximum or minimum of the given options, it suggested that stronger or weaker penalties, respectively, might be needed. While we did not explore values beyond these boundaries to avoid protracted computational effort, we highlight in the appendix how often users might encounter this issue under varying sample sizes and should consider expanding the range of penalty values explored. Specifically, we report the frequency of penalty estimates at the extremes of selected values in our network and covariate complexity scenarios (see *Appendix S1: Figure S5*).

We assessed model performance based on percent relative bias 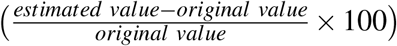 of the natural parameters (*f* ‘s), general parameters (*ψ*), species-specific marginal probabilities (e.g., *ψ*_*A*_), and conditional occupancy probabilities (e.g. *Pr*(*Z*_*A*_ = 1|*Z*_*B*_ = 1)). We also calculated the power (i.e. inverse false negative rate) to detect a statistically significant coefficient for all parameters (intercept and slopes) within second-order natural parameters (e.g. *f*_12_ = *γ*_0_ *≠*0) under a convention of type-I error rate of *α <* 0.05. We considered bias below 10% to be negligible and an acceptable power of ≥80%. We calculated performance metrics using only models that both converged and had identifiable parameters that were not boundary estimates, defined as those whose absolute value is less than *logit*(0.99992) = 9.3 ((Welsh et al. 2013); see Appendix S1:Table S2 for a summary of the frequency of convergence issues). All code for simulating data and model fitting is available at Cowans et al. (2025).

## Results

Here, we present the power of the co-occurrence model to estimate the interaction coefficient (*f*_12_) and the parameter bias under varying sample sizes for each of our simulation scenarios. We refer to an unbiased estimate as one with a percent relative bias below 10%. In all tables, we denote >10% bias in bold, italise biases between 5-10% and replace bias values <5% with dots (“.”).

### Null models for two interacting species

We encountered no convergence issues in the null model scenarios and therefore only report results using the log-likelihood approach, noting the general patterns of power and bias across sample size using penalized likelihood were consistent. We observed a clear effect of sample size, interaction strength, and detection probability on the power to detect and estimate the correct interaction coefficient (Figure 1). Under this simplest model parameterization, with high detection probability, estimation of the interaction parameter with 80% power was reached at around 200 sites with strong interactions and 600 sites with moderate (Figure 1). Power to detect interactions decreased with low detection probability; 80% power was reached at around 600 and 900 sites with strong negative and positive interactions, respectively, and moderate interactions required around 2300 sites to reach nominal power (Figure 1). At the maximum sample size, the power to detect weak interactions was approximately 50% under high detection probability and was less than 25% when detection probability was low.

**Figure 1:**
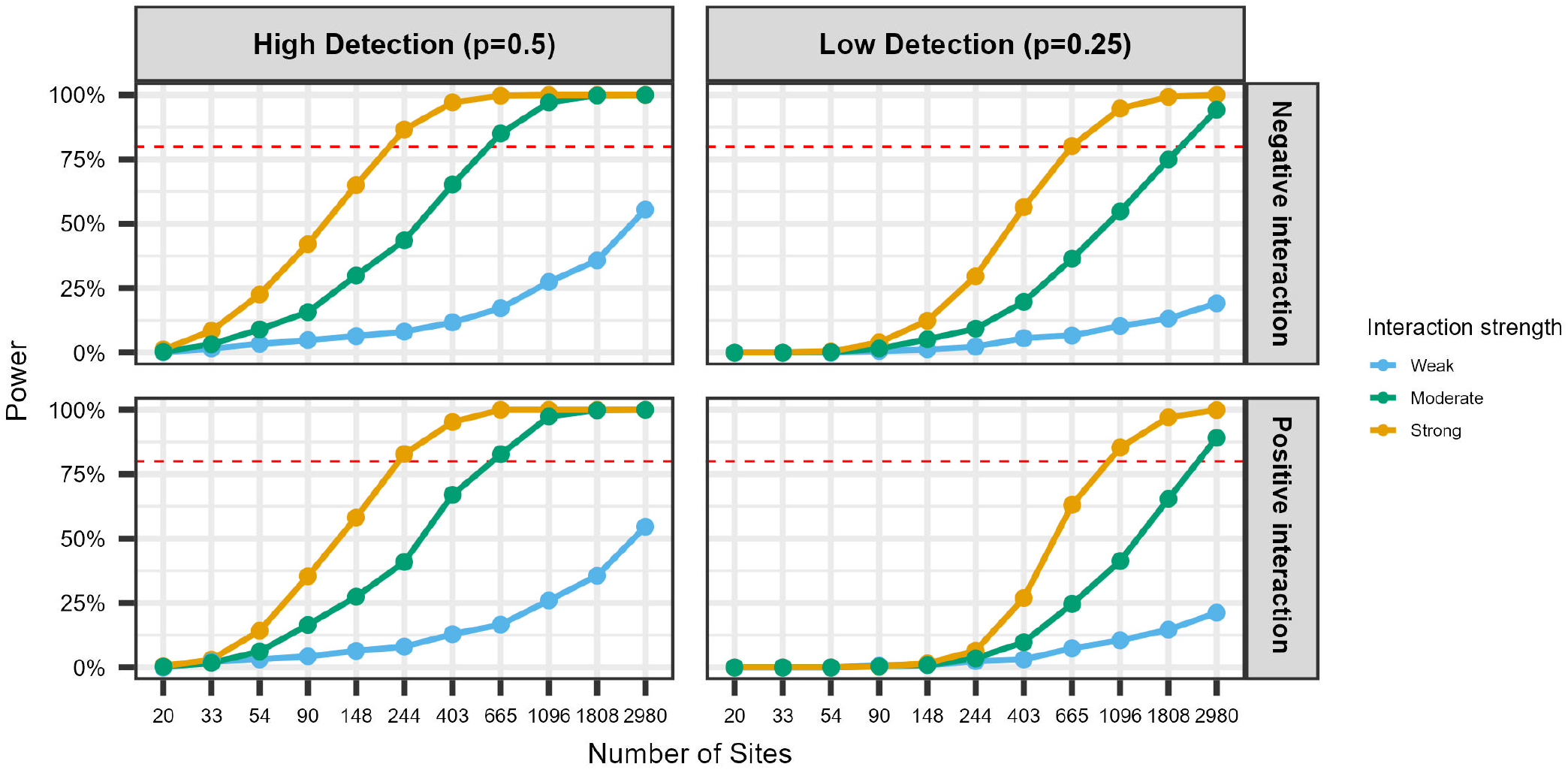
Power (%) of the null two species co-occurrence model to detect strong (yellow; *γ*_0_ = *±*1), moderate (green; *γ*_0_ = *±*0.6) and weak (blue; *γ*_0_ = *±*0.2) significant interactions under increasing sample sizes. A power of 80% is shown by the horizontal red line.

Unbiased estimates (i.e., percent relative bias under 10%) of the interaction parameter (*f*_12_) were achieved with 90 sites for strong negative interactions and 148 sites for weak negative interactions under high detection probability (Table 2). When detection probability was low, this number increased to 403 and 1096 sites, respectively (Table 2). Unbiased estimation of first-order natural parameters (*f*_1_, *f*_2_) required at least 665 sites when detection was high and between 1808 and 2980 sites when detection was low (Table 2). General parameters (*ψ*) were less sensitive: reliable estimates were obtained at 244 sites for low detection scenarios and 33 sites for high detection (Table 2). Marginal and conditional occupancy probabilities were similarly unbiased, requiring as few as 20 or 33 sites for high detection (for weak and strong interactions, respectively), and 54 or 148 sites for low detection (Table 3). Results for parameter bias with strong and weak positive interactions can be found in the Appendix (Appendix S1: Table S3). Notably, while interaction sign had little effect under high detection, positive interactions were associated with reduced power and increased bias under low detection, requiring larger sample sizes than negative interactions to achieve comparable performance (Appendix S1: Table S3). This pattern may reflect how negative interactions reduce the chance to observe both species co-occurring and thus the ability to correctly estimate the data-generating values. However, we caution that these patterns may be partly artefactual, influenced by specific data-generating values or stochastic variation in simulation and model fitting.

**Table 2:** Mean relative bias (%RB) of natural (f) and general (ψ) parameters for the two species null model with weak and strong negative interactions. We denote >10% bias in bold, italise biases between 5-10% and replace bias values <5% with dots (“.”).

| N | Strength | High Detection |  |  |  |  |  |  | Low Detection |  |  |  |  |  |  |
| --- | --- | --- | --- | --- | --- | --- | --- | --- | --- | --- | --- | --- | --- | --- | --- |
|  |  | Natural |  |  | General |  |  |  | Natural |  |  | General |  |  |  |
| | | $f_1$ | $f_2$ | $f_{12}$ | $\psi_{00}$ | $\psi_{10}$ | $\psi_{01}$ | $\psi_{11}$ | $f_1$ | $f_2$ | $f_{12}$ | $\psi_{00}$ | $\psi_{10}$ | $\psi_{01}$ | $\psi_{11}$ |
| 20 | Weak | **281.27** | **398.45** | **85.91** | **-11.24** | . | . | 8.33 | **578.77** | **664.75** | **-389.86** | **-18.58** | -9.34 | -9.05 | **38.9** |
| 33 |  | **202.46** | **148.33** | **36.72** | -7.45 | . | . | 7.39 | **709.35** | **851.58** | **-32.23** | **-18.99** | -5.34 | . | **26.25** |
| 54 |  | **122.94** | **110.23** | **49.12** | -5.48 | . | . | . | **631.11** | **665.09** | **59.22** | **-17.27** | . | . | **18.01** |
| 90 |  | **40.81** | **50.72** | **20.32** | . | . | . | . | **406.72** | **270.06** | **39.11** | **-10.7** | . | . | **10.91** |
| 148 |  | 9.47 | **11.8** | . | . | . | . | . | **162.42** | **195.33** | . | -8.13 | . | . | **10.93** |
| 244 |  | **15.12** | **10.18** | . | . | . | . | . | **131.42** | **124.48** | **35.66** | -6.73 | . | . | 5.89 |
| 403 |  | **11.88** | . | . | . | . | . | . | **74.75** | **45.85** | **19.43** | . | . | . | . |
| 665 |  | . | 6.13 | . | . | . | . | . | **44.77** | **46.45** | **14.47** | . | . | . | . |
| 1096 |  | . | 9.25 | . | . | . | . | . | **16.33** | **21.44** | 5.94 | . | . | . | . |
| 1808 |  | . | . | . | . | . | . | . | **10.66** | 7.49 | . | . | . | . | . |
| 2980 |  | . | . | . | . | . | . | . | . | . | . | . | . | . | . |
| 20 | Strong | **488.93** | **412.47** | **40.61** | **-12.8** | . | . | **24.45** | **649.27** | **788.84** | -5.48 | **-16.19** | **-21.37** | **-11.85** | **117.72** |
| 33 |  | **232.17** | **263.18** | **28.1** | -8.6 | . | . | 9.3 | **974.22** | **992.35** | **61.63** | **-19.2** | -5.26 | . | **66.48** |
| 54 |  | **79.53** | **102.41** | **14.11** | . | . | . | . | **886.38** | **897.96** | **67.53** | **-20.07** | . | . | **41.93** |
| 90 |  | **40.51** | **57.73** | 7.84 | . | . | . | . | **843.57** | **777.29** | **72.56** | **-19.52** | 6.51 | . | **21.48** |
| 148 |  | **21.49** | **33.72** | . | . | . | . | . | **400.5** | **387.49** | **34.07** | **-12.17** | . | . | **15.24** |
| 244 |  | **14.63** | **13.3** | . | . | . | . | . | **179.6** | **182.42** | **17.21** | -8.16 | . | . | 7.64 |
| 403 |  | **20.3** | **14.94** | . | . | . | . | . | **95.86** | **87.4** | 9.58 | . | . | . | . |
| 665 |  | . | . | . | . | . | . | . | **54.6** | **54.41** | 5.27 | . | . | . | . |
| 1096 |  | . | . | . | . | . | . | . | **29.6** | **21.15** | . | . | . | . | . |
| 1808 |  | . | . | . | . | . | . | . | **17.67** | **20.87** | . | . | . | . | . |
| 2980 |  | . | . | . | . | . | . | . | 8.11 | 8.03 | . | . | . | . | . |

**Table 3:** Mean relative bias (%RB) of marginal and conditional probability for the two species null model with weak and strong negative interactions. We denote >10% bias in bold, italise biases between 5-10% and replace bias values <5% with dots (“.”).

| N | Strength | High Detection |  |  |  |  |  | Low Detection |  |  |  |  |  |
| --- | --- | --- | --- | --- | --- | --- | --- | --- | --- | --- | --- | --- | --- |
|  |  | Marginal |  | Conditional |  |  |  | Marginal |  | Conditional |  |  |  |
|  |  | Sp1 | Sp2 | Sp1=1\|Sp2=0 | Sp1=1\|Sp2=1 | Sp2=1\|Sp1=0 | Sp2=1\|Sp1=1 | Sp1 | Sp2 | Sp1=1\|Sp2=0 | Sp1=1\|Sp2=1 | Sp2=1\|Sp1=0 | Sp2=1\|Sp1=1 |
| 20 | Weak | . | 5.65 | 5.04 | . | 7.2 | 5.13 | **13.58** | **13.73** | 5.47 | **21.18** | . | **21.86** |
| 33 |  | . | . | . | . | . | . | 9.66 | **11.83** | 7.51 | **10.86** | **10.37** | **13.57** |
| 54 |  | . | . | . | . | . | . | 8.12 | 8.64 | 7.35 | 8.93 | 8.94 | 9.65 |
| 90 |  | . | . | . | . | . | . | 5.84 | . | 6.24 | 6.81 | . | . |
| 148 |  | . | . | . | . | . | . | . | . | . | 5.9 | . | 6.41 |
| 244 |  | . | . | . | . | . | . | . | . | . | . | . | . |
| 20 | Strong | 9.24 | 6.06 | 9.16 | **20.63** | 6.66 | **16.17** | **18.84** | **25.61** | . | **61.35** | . | **68.67** |
| 33 |  | . | 5.03 | 5.18 | . | 7 | 6.55 | **15.47** | **16.1** | **11.48** | **39.53** | **11.13** | **40.22** |
| 54 |  | . | . | . | . | . | . | **12.41** | **12.62** | **13.38** | **22.86** | **13.86** | **23.47** |
| 90 |  | . | . | . | . | . | . | **10.84** | 7.93 | **16.38** | **12.89** | **14.09** | 8.91 |
| 148 |  | . | . | . | . | . | . | 6.55 | 5.68 | 9.48 | 9.07 | 8.75 | 8.14 |
| 244 |  | . | . | . | . | . | . | . | . | 6.09 | . | 5.81 | . |
| 403 |  | . | . | . | . | . | . | . | . | . | . | . | . |

### Null models for more than two species: network complexity

Models that included more species frequently experienced convergence issues at low sample sizes. Therefore, we report results only from simulations with more than 50 sites and summarize convergence issues in the Appendix (Appendix S1: Table S2). As with the simpler model, power and bias were primarily influenced by sample size and interaction strength. Increasing the number of species reduced the power to detect interactions of similar strength (i.e., comparing lines of the same color as species count increased from 3 to 5; Figure 2). This reduction in power was observed in both likelihood methods until the number of sites exceeded 1,096, and was more pronounced in low sample sizes for the log-likelihood approach. For the penalized likelihood, the reduction was most evident at moderate interaction strengths and smaller sample sizes (Figure 2). When using the log-likelihood method, 400 sites were required to estimate strong interactions with 80% power for a model with 5 species compared to around 250 sites when only 3 species were included (Figure 2). Below 400 sites, penalized likelihood returned greater power than log-likelihood scenarios irrespective of number of species and interaction strength, highlighting the benefits of this approach under small sample sizes.

**Figure 2:**
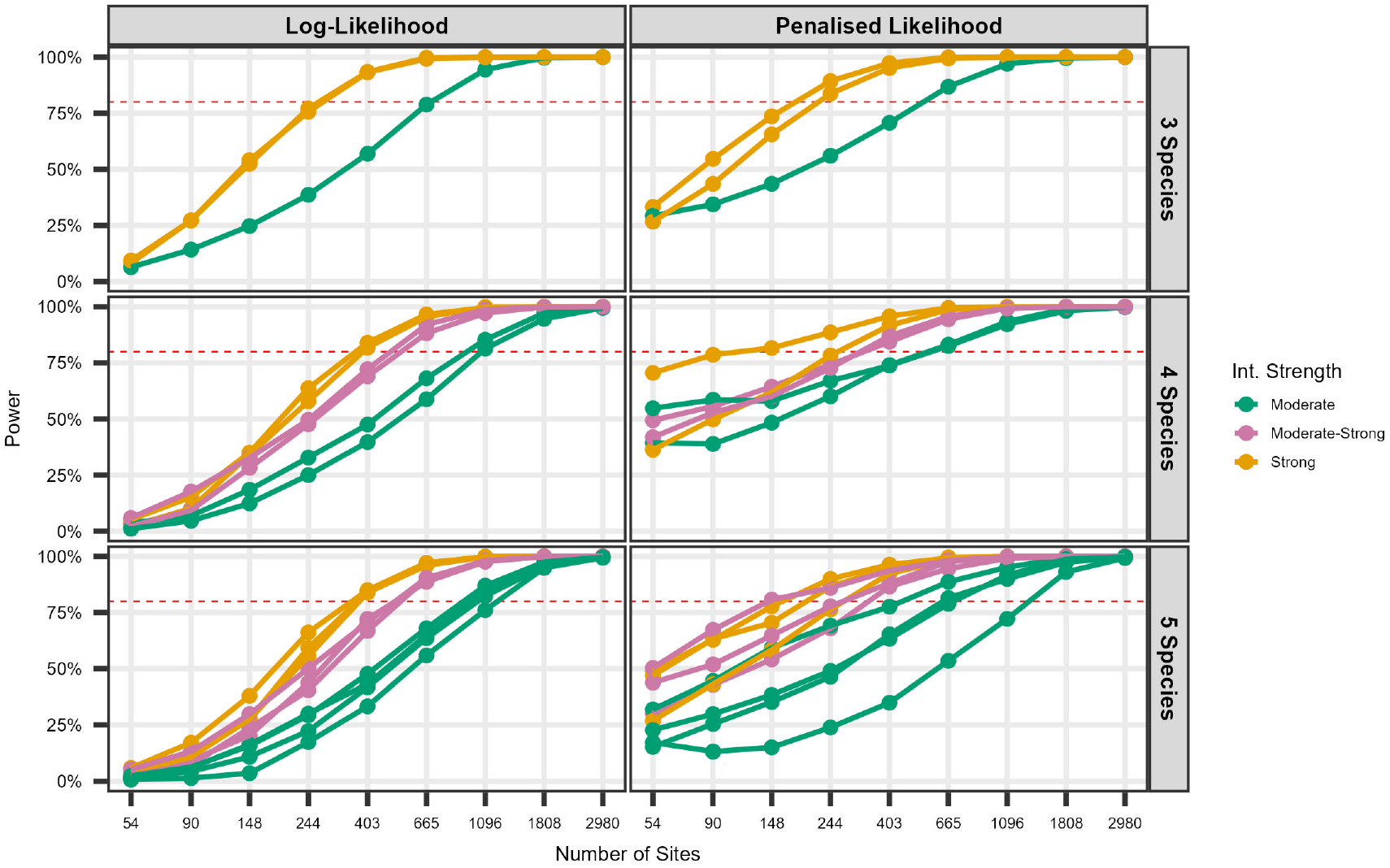
Power of the co-occurrence model to detect a significant interaction term with increasing network complexity (3-5 species). Each line represents one pairwise interaction term (e.g. with 4 species there are 6 pairwise terms, with 5 species there are 10 pairwise interaction terms) and each term is colored by the strength.

Log-likelihood estimates of all pairwise natural parameters eventually converged on minimal relative bias, requiring 403 sites in the 5 species model (Table 4). As expected, penalized likelihood estimates of natural parameters continued to show bias in the highest sample size scenario (Table 4). Extreme penalty values could modulate bias and while frequent at the smallest sample sizes, they ceased to occur beyond *N >* 403 (Appendix S1: Figure S5). For the log-likelihood method, general parameters were unbiased at 54 sites for 3 species, 90 with 4 species and 148 for 5 species (Appendix S1: Figure S2). Estimates of marginal occupancy probabilities were unbiased across all sample sizes above 50 for models including 3, 4 and 5 species under both log and penalized likelihood (Appendix S1: Figure S2), although unbiased penalized likelihood estimates of conditional occupancy probabilities were not reached until 1808 sites (Appendix S1: Figure S3).

**Table 4:** Mean relative bias (%RB) of second-order natural (f) parameters in the scenario of 5 interacting species under normal (LL) and penalized (PL) likelihood estimation methods. Natural parameter bias results for 3 and 4 species can be visualised in Appendix S1: Figure S3. We denote >10% bias in bold, italise biases between 5-10% and replace bias values <5% with dots (“.”).

| N | Likelihood | $f_{12}$ | $f_{13}$ | $f_{14}$ | $f_{15}$ | $f_{23}$ | $f_{24}$ | $f_{25}$ | $f_{34}$ | $f_{35}$ | $f_{45}$ |
| --- | --- | --- | --- | --- | --- | --- | --- | --- | --- | --- | --- |
| **54** | LL | **32.37** | -6.99 | **28.85** | **13.85** | **27.98** | **10.32** | **36.92** | **25.06** | 5.7 | **12.73** |
| **90** |  | **24.39** | **18.84** | **23.61** | **36.42** | **19.17** | 8.79 | **22.78** | **18.89** | **20.1** | **26.32** |
| **148** |  | **11.11** | **19.89** | **12.46** | **24.65** | **12.22** | . | **12.99** | 6.42 | **16.62** | **16.14** |
| **244** |  | 9.45 | 5.64 | 8.34 | **14.19** | 5.78 | . | 8.5 | . | . | **11.38** |
| **403** |  | 5.05 | 7.29 | 8.17 | 9.06 | . | . | 5.1 | . | . | 6.64 |
| **665** |  | . | . | . | 5.36 | . | . | . | . | . | . |
| **1096** |  | . | . | . | . | . | . | . | . | . | . |
| **54** | PL | **-47.77** | **-53.96** | **-50.31** | **-71.47** | **-44.4** | **-30.43** | **-59.17** | **-38.2** | **-30.45** | **-47.87** |
| **90** |  | **-32.82** | **-33.9** | **-33.87** | **-63.1** | **-29.84** | -9.5 | **-44.28** | **-22.67** | -9.45 | **-29.63** |
| **148** |  | **-27.21** | **-24.43** | **-25.54** | **-58.26** | **-22.39** | . | **-37.59** | **-16.77** | . | **-21.67** |
| **244** |  | **-18.38** | **-17.18** | **-18.7** | **-46.82** | **-15.68** | 6.93 | **-28.4** | **-10.15** | . | **-14.82** |
| **403** |  | **-14.3** | -8.51 | **-12.13** | **-37.57** | **-12.61** | 5.27 | **-22.05** | -8.55 | . | **-11.7** |
| **665** |  | **-10.47** | -6.38 | **-12.68** | **-29.17** | -8.69 | 5.57 | **-17.57** | -5.2 | . | -9.71 |
| **1096** |  | -8.52 | . | -8.03 | **-22.92** | -6.67 | . | **-12.75** | . | . | -7.74 |
| **1808** |  | -5.12 | . | -5.98 | **-14.51** | . | . | -8.14 | . | . | -5.88 |
| **2980** |  | . | . | . | **-10.58** | . | . | -5.55 | . | . | . |

### Covariate models for two species: covariate complexity

Regardless of effect size and model structure, statistical power was generally higher and reached nominal levels more rapidly for slope parameters compared to intercepts (Figure 3), with the latter exhibiting trends similar to those observed in the null models. More complex model structures required higher sample sizes relative to simpler covariate structures to achieve nominal bias levels (Table 5). Under log-likelihood, unbiased estimates were obtained at 90 sites with one unique covariate, 148 sites with one shared, and 244 sites with one unique and one shared (Table 5). As in the network scenarios, the advantages of penalized likelihood in increasing power were most evident with fewer than 200 sites, particularly when effect sizes were moderate-to-strong or strong (Figure 3), but likely due to shrinkage,estimates remained negatively biased beyond *N >* 400 (Table 4) where extreme penalty values are no longer observed (Figure S5).

**Table 5:** Mean relative bias (%RB) of the coefficient regression parameters (γ_0_, γ_1_, γ_2_) in the three covariate complexity scenarios (Unique, Shared, Unique+Shared) fitted under normal (LL) and penalized (PL) likelihood methods. We denote >10% bias in bold, italise biases between 5-10% and replace bias values <5% with dots (“.”).

| N | Likelihood | Unique |  | Shared |  | Unique + Shared |  |  |
| --- | --- | --- | --- | --- | --- | --- | --- | --- |
| | | $\gamma_0 = 0.8$ | $\gamma_1 = -1$ | $\gamma_0 = 0.6$ | $\gamma_1 = 0.6$ | $\gamma_0 = 0.6$ | $\gamma_1 = -1$ | $\gamma_2 = 0.8$ |
| **54** | LL | **18.83** | **21.76** | **34** | **30.03** | . | **31.67** | **54.8** |
| **90** |  | 5.57 | 9.58 | **12.81** | **12.02** | . | **18.84** | **20.36** |
| **148** |  | . | 5.05 | 6.36 | 5.31 | . | 8.86 | **11.42** |
| **244** |  | . | . | . | . | . | . | . |
| **54** | PL | **-57.48** | **-30.72** | **-70.33** | **-57.01** | **-62.29** | **-33.38** | **-26.2** |
| **90** |  | **-48.41** | **-21.94** | **-65.69** | **-47.39** | **-50.25** | **-21.96** | **-22.22** |
| **148** |  | **-36.01** | **-13.92** | **-49.03** | **-33.62** | **-37.67** | **-16.13** | **-16.06** |
| **244** |  | **-23.78** | -9.41 | **-36.11** | **-23.14** | **-24.89** | **-11.51** | **-11.75** |
| **403** |  | **-16.77** | . | **-22.52** | **-12.9** | **-18.86** | -7.06 | -5.99 |
| **665** |  | -9.58 | . | **-13.32** | -7.79 | **-13.77** | . | -5.76 |
| **1096** |  | -5.03 | . | -8.87 | -5.76 | -8.47 | . | . |
| **1808** |  | . | . | . | . | -6.7 | . | . |
| **2980** |  | . | . | . | . | . | . | . |

**Table 6:** Recommended minimum sample sizes for the Rota et al. (2016) co-occurrence model based on inference objective. The values represent the number of sites required to estimate the interaction term with both 80% power and less than 10% bias (column 1) or estimate marginal/-conditional occupancy probabilities with less than 10% bias (column 2). The exact number of sites will depend on specific parameter values, and what we report is based on the patterns we observed within our simulation sets involving null models for two interacting species.

| Scenario | Recommended minimum number of sites |  |
| --- | --- | --- |
| | Identify interactions via $f_{12}$ | Generate unbiased occupancy predictions |
| High detection $p^* = 0.88$<br>Strong interactions $\gamma_0 = 1$ | 244 | 33 |
| Low detection $p^* = 0.58$<br>Strong interactions $\gamma_0 = 1$ | 665 ($f_{12} < 0$)<br>1096 ($f_{12} > 0$) | 148 |
| High detection $p^* = 0.88$<br>Weak interactions $\gamma_0 = 0.2$ | Greater sample size than examined | 20 |
| Low detection $p^* = 0.58$<br>Weak interactions $\gamma_0 = 0.2$ | Greater sample size than examined | 54 |

**Figure 3:**
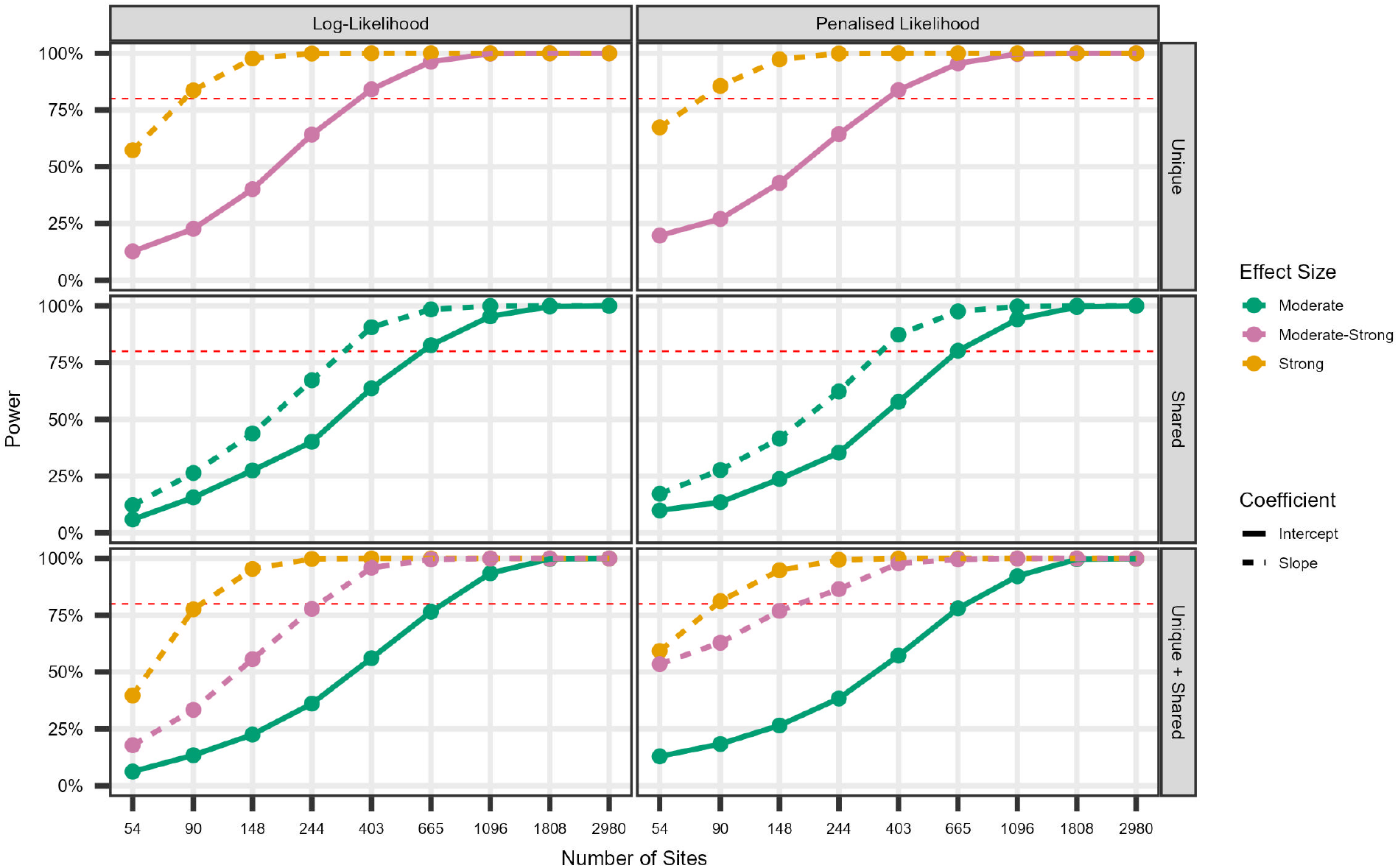
Relationship between the Power (%) and sample size for the intercept (solid) and slope (dashed) regression coefficients (*γ*_0_, *γ*_1_, *γ*_2_) in the three covariate complexity scenarios (*Unique, Shared, Unique+Shared*) fitted under normal (LL) and penalized (PL) likelihood methods. Effect size varied between moderate (*γ* =±0.6; green), moderate-strong (*γ* =±0.8; pink) and strong (*γ* =±1; orange).

## Discussion

The rise of accessible remote sensing technologies that simultaneously monitor multiple species, such as camera traps and autonomous recording units, has significantly advanced the study of species co-occurrence and its determinants using large-scale observational data (Burton et al. 2015; Kerry et al. 2022). This progress has facilitated the widespread use of co-occurrence models, designed to detect inter-specific dependencies in occurrence over and above habitat effects (Kéry and Royle 2020; Devarajan et al. 2020). Although it is widely reported that these dependencies are statistical associations (Rota et al. 2016; Blanchet et al. 2020), they may reflect underlying biological interactions (Twining et al. 2022; Parsons et al. 2019), though the connection between statistical co-occurrence and ecological interaction remains complex and non-trivial (Thurman et al. 2019; Freilich et al. 2018).

By evaluating the performance of a widely applied co-occurrence model across a range of biologically realistic scenarios, we identify several key considerations when drawing inferences about ecological processes. Our simulations show that, under the simplest model structure, the number of sites required to detect species interactions is substantial and varies according to detection probability and interaction strength. Strong interactions typically require 200–1000 sites, depending on species detectability, while weak interactions demand far larger sample sizes and are likely infeasible to estimate even under high detectability.

We also show that while sample sizes below c.150 consistently limited the ability to infer interaction strength, unbiased estimates of general parameters and marginal and conditional occupancy probability required fewer sites. Natural parameters (*f*_1_, *f*_2_, *f*_12_), which underpin inference on ecological interactions, are highly biased at low sample sizes, especially when detection is poor. In contrast, general parameters (***ψ***) and marginal and conditional occupancy probabilities, are estimated with minimal bias with as few as 50 sites. This discrepancy raises an important consideration for both model use and interpretation. While general parameters may appear more robust and therefore preferable in applied contexts, they do not directly inform the ecological interactions between species - the very relationships multispecies co-occurrence models are often used to investigate. If only general parameters are reliably estimated under realistic sample sizes, then the primary utility of these models may lie in identifying patterns of co-occurrence rather than inferring underlying processes. As noted in Rota et al. (2016), such models are best suited for hypothesis generation, providing insight into, but not definitive evidence of, the mechanisms driving observed patterns. This distinction is important for guiding interpretation and setting appropriate expectations for inference from co-occurrence data.

A critical insight of this study is that reliable ecological inference using multispecies occupancy models depends not only on the choice of model structure but on clarity about inferential goals (Tredennick et al. 2021). If the aim is to infer co-occurrence presence and/or strength, researchers must plan for large sample sizes and high detection probabilities. Conversely, if the goal is to predict site occupancy patterns across a landscape, general parameters and derived occupancy probabilities may suffice with more modest data requirements (Table 5). This highlights the need for careful alignment between model goals, data collection strategy, and the specific parameters targeted for inference.

When using these models to predict occupancy probabilities, penalized likelihood fitting frameworks have already been developed (Clipp et al. 2021). In our study, we explicitly demonstrate how the bias–variance trade-off inherent in penalized likelihood estimation varies across sample sizes, a feature we feel is underappreciated. In particular, we show that penalized likelihood offers clear benefits at sample sizes below 400 sites, including improved power and convergence and reduced variance, resulting in more stable estimates, albeit with increased bias. Extreme penalty values when using default optimization settings occurred the most frequently at sample sizes under 400 sites. When using default settings, users can expect extreme penalties to occur 10–60% of the time, depending on model structure, for sample sizes below 100 (Appendix S1: Figure S5), and we recommend that users examine the 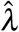 estimates and adjust values outside the default range in these scenarios.

Our findings are consistent with recent work showing parameter precision increases with the strength of species avoidance, and that the number of sites has a greater impact on precision than temporal replication (Louvrier et al. 2024). In simulations applying a two-species null model, the co-occurrence model only outperformed a non-interaction model when avoidance was strong and at least 150 sites were sampled (Louvrier et al. 2024). Collectively, these findings underscore a persistent challenge in ecological network inference: reliably detecting weak or moderate interactions remains difficult, particularly when datasets are small.

In our simulations, increasing model complexity through the number of interacting species and covariates increased sample size requirements to achieve acceptable power and bias. However reliable inference was overall far more sensitive to the number of sites and the strength of the interaction of interest than to model complexity. We note that we considered only second-order interactions here, but feel this is appropriate given this is the most commonly used approach (Kéry and Royle 2020). This suggests that, when sample sizes are sufficiently large, more complex models can reliably detect strong interactions between multiple species’ occupancy probabilities. However, detecting weaker associations consistently requires high sample sizes regardless of model complexity. Even under its simplest formulation and high detection probabilities, the sample size requirements for weak interactions exceeded that of any study published using this model (Figure S1). Our results therefore provide compelling evidence that likely many studies inferring no interactions may instead reflect data limitations rather than a true lack of association, echoing broader concerns in the general application of co-occurrence models (Blanchet et al. 2020).

Given this, we emphasize the need for further methodological development that improves inference when sample sizes are constrained and interactions are moderate. Promising directions include Bayesian frameworks with informative priors, which can mitigate boundary estimate issues in similar contexts (Fidino et al. 2019), and integrated modelling strategies that leverage multiple data streams to improve convergence issues and inference ability in small sample size cases (Lauret et al. 2023). For example, camera trap data can be integrated with point counts or transect surveys to provide a more comprehensive understanding of species distributions and co-occurrence patterns (Lauret et al. 2023). We note integrated models will not circumvent high sample size requirements but will allow more information to be accumulated through combining multiple datasets. While our study assumed three sampling occasions (J = 3), future work may also explore the effect of increased temporal replication on required sample sizes when cumulative detection probabilities remain below one (Guillera-Arroita et al. 2010). However, existing evidence suggests additional temporal effort alone is insufficient to reduce the sample size needed to reliably detect species interactions in null models (Louvrier et al. 2024).

Despite wide application across a range of sampling scenarios from e.g. 40 sites (Carricondo-Sanchez et al. 2019) to over 2000 (Mandeville et al. 2024), practical guidance on fitting multispecies models, including considerations for model selection and interpretation, are still lacking. Our results support the general principle that increasing the number of sites improves inference performance, even in more complex ecological networks. However, we acknowledge that real-world ecological interactions are spatio-temporally dynamic. This variability can make inference more challenging and, in some cases, require an impractically large number of sites to achieve reliable estimates. As such, while larger sample sizes generally enhance model performance, the feasibility of this approach depends on the ecological context. Rather than prescribing universal guidelines, we emphasise the need for careful consideration of study-specific factors, including temporal and spatial variation in interactions, when interpreting co-occurrence patterns.

In this study, we provide an important context for linking observational data and ecological inference about biotic and abiotic determinants of species distributions. Specifically, we provide key sample size considerations in a range of contexts to allow reliable estimation of a non-independence in species occurrence states. Only then, researchers can consider what these associations may reflect in terms of biological interactions between species, on which an emerging literature exists (Blanchet et al. 2020; Thurman et al. 2019; Freilich et al. 2018). Although the Rota et al. (2016) model is notable for allowing associations to vary as a function of covariates, other co-occurrence frameworks such as the species interaction factor (SIF) model of MacKenzie et al. (2004) have largely fallen out of use in multispecies applications due to high sample size requirements (Kerry et al. 2022), and we believe it would be interesting to revisit SIFs with the renewed sample sizes considered here in mind. Our findings suggest that high sample sizes are not specific to any one framework but are instead a general feature of models that attempt to estimate species interactions under imperfect detection. These requirements should not be viewed as shortcomings of the models themselves, but rather as a reflection of the limited information content in typical observational ecological datasets. In multispecies occupancy models, data are inherently noisy and sparse, especially with regard to the joint detection histories required to estimate interaction parameters. As such, it is not unexpected that robust inference of complex ecological processes like species interactions necessitates large sample sizes. Sophisticated modeling approaches cannot compensate for fundamentally low-information data, where reliable inference in ecology always demands adequate sampling effort (Warton et al. 2015).

## Acknowledgements

AC, ABB, and CS conceptualized the study, conducted simulations, and wrote and edited the manuscript.

