## Appendix S1 for "Sample size considerations for species co-occurrence models"

This supplement contains additional tables and figures for the simulation scenarios described in "Sample size considerations for species co-occurrence models."

### Literature review

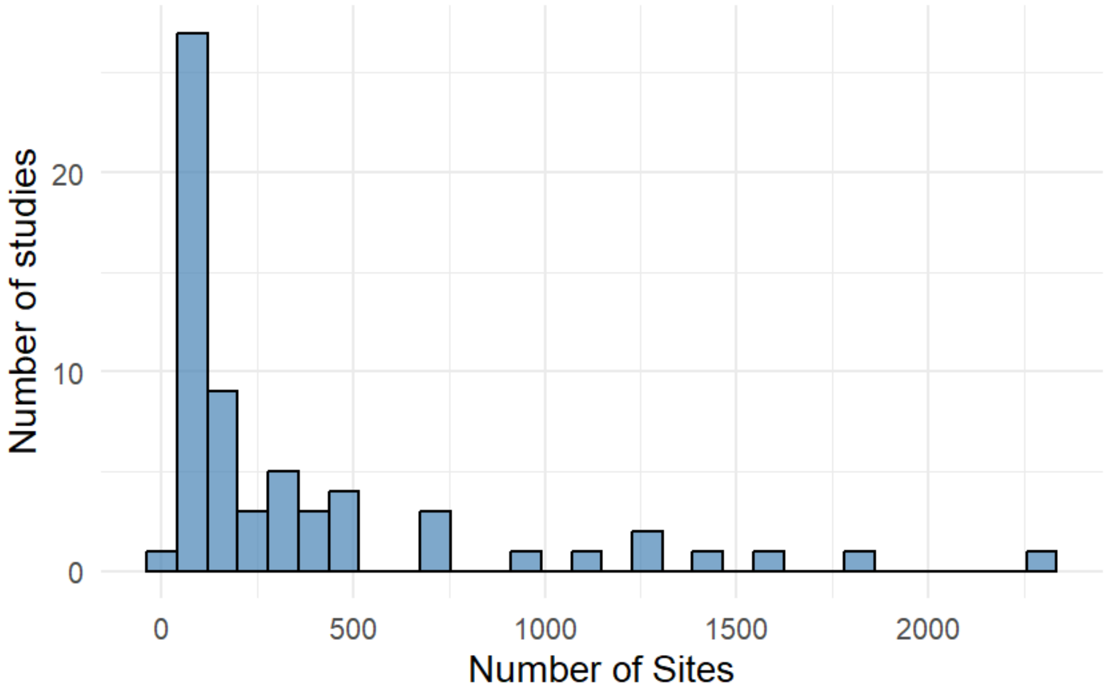


Figure S1: *Distribution of sample sizes applying the multispecies occupancy model of Rota et al. (2016) as of April 2025.*

### Data Generating Values

Table S1: *Table containing the data generating parameter values for all 18 scenarios, split into first order (single species occupancy log-odds) and second order (pairwise interactions) natural parameters.*

| | Scenario | First order | Second order |
|---|---|---|---|
| Null High detection | Weak+ | $f_1 = f_2 = -0.4$ | $f_{12} = \gamma_0 = 0.2$ |
| | Moderate+ | $f_1 = f_2 = -0.2$ | $f_{12} = \gamma_0 = 0.6$ |
| | Strong+ | $f_1 = f_2 = -0.4$ | $f_{12} = \gamma_0 = 1$ |
| | Weak- | $f_1 = f_2 = 0.1$ | $f_{12} = \gamma_0 = -0.2$ |
| | Moderate- | $f_1 = f_2 = 0.2$ | $f_{12} = \gamma_0 = -0.6$ |
| | Strong- | $f_1 = f_2 = 0.1$ | $f_{12} = \gamma_0 = -1$ |
| Null Low Detection | Weak+ | $f_1 = f_2 = -0.4$ | $f_{12} = \gamma_0 = 0.2$ |
| | Moderate+ | $f_1 = f_2 = -0.2$ | $f_{12} = \gamma_0 = 0.6$ |
| | Strong+ | $f_1 = f_2 = -0.4$ | $f_{12} = \gamma_0 = 1$ |
| | Weak- | $f_1 = f_2 = 0.1$ | $f_{12} = \gamma_0 = -0.2$ |
| | Moderate- | $f_1 = f_2 = 0.2$ | $f_{12} = \gamma_0 = -0.6$ |
| | Strong- | $f_1 = f_2 = 0.1$ | $f_{12} = \gamma_0 = -1$ |
| Covariate | Unique | $f_1 = -0.4 + 0.6X_A; f_2 = -0.6 + 0.8X_B$ | $f_{12} = 0.8 - 1X_C$ |
| | Shared | $f_1 = -0.4 + 0.6X_A; f_2 = -0.6 + 0.8X_B$ | $f_{12} = 0.6 - 0.6X_A$ |
| | Unique+Shared | $f_1 = -0.4 + 0.6X_A; f_2 = -0.6 + 0.8X_B$ | $f_{12} = 0.6 - 1X_C + 0.8X_A$ |
| Network | 3 Species | $f_1 = -0.4; f_2 = 0.3; f_3 = 0.8$ | $f_{12} = 1; f_{13} = -0.6; f_{23} = -1$ |
| | 4 Species | $f_4 = -0.2$ | $f_{14} = 0.8; f_{24} = 0.6; f_{34} = -0.8$ |
| | 5 Species | $f_5 = 0.5$ | $f_{15} = 0.6; f_{25} = -1; f_{35} = 0.8; f_{45} = -0.6$ |

#### Convergence Rates

Table S2: *Convergence rates under log likelihood (LL) and penalized likelihood (PL) estimation methods for the simplest (Null) and increased complexity (Network and Covariate) models. All Null models (except one in scenario 6) converged under LL, whereas increasing complexity reduced convergence rates, dropping below 50% in the scenario with 5 species at the smallest sample size. All scenarios using PL converged except 3 cases in Network and 2 in Covariate models, showcasing the benefit of the PL framework in improving convergence for more complex models at small sample sizes.*

| | | Null models | | | | | | Network models | | | Covariate models | | |
|---|---|---|---|---|---|---|---|---|---|---|---|---|---|
| N | Likelihood | 1 | 2 | 3 | 4 | 5 | 6 | 3Spp. | 4Spp. | 5Spp. | Unique | Shared | Unique+Shared |
| **20** | LL | 1.00 | 1.00 | 1.00 | 1.00 | 1.00 | 0.99 | 0.93 | 0.79 | 0.44 | 0.91 | 0.96 | 0.85 |
| **33** | LL | 1.00 | 1.00 | 1.00 | 1.00 | 1.00 | 1.00 | 0.94 | 0.91 | 0.66 | 0.96 | 0.97 | 0.91 |
| **54** | LL | 1.00 | 1.00 | 1.00 | 1.00 | 1.00 | 1.00 | 0.96 | 0.95 | 0.73 | 0.98 | 0.98 | 0.96 |
| **90** | LL | 1.00 | 1.00 | 1.00 | 1.00 | 1.00 | 1.00 | 0.99 | 0.97 | 0.86 | 0.99 | 1.00 | 0.99 |
| **148** | LL | 1.00 | 1.00 | 1.00 | 1.00 | 1.00 | 1.00 | 1.00 | 1.00 | 0.94 | 1.00 | 1.00 | 1.00 |
| **244** | LL | 1.00 | 1.00 | 1.00 | 1.00 | 1.00 | 1.00 | 1.00 | 1.00 | 0.99 | 1.00 | 1.00 | 1.00 |
| **403** | LL | 1.00 | 1.00 | 1.00 | 1.00 | 1.00 | 1.00 | 1.00 | 1.00 | 1.00 | 1.00 | 1.00 | 1.00 |
| **665** | LL | 1.00 | 1.00 | 1.00 | 1.00 | 1.00 | 1.00 | 1.00 | 1.00 | 1.00 | 1.00 | 1.00 | 1.00 |
| **1096** | LL | 1.00 | 1.00 | 1.00 | 1.00 | 1.00 | 1.00 | 1.00 | 1.00 | 1.00 | 1.00 | 1.00 | 1.00 |
| **1808** | LL | 1.00 | 1.00 | 1.00 | 1.00 | 1.00 | 1.00 | 1.00 | 1.00 | 1.00 | 1.00 | 1.00 | 1.00 |
| **2980** | LL | 1.00 | 1.00 | 1.00 | 1.00 | 1.00 | 1.00 | 1.00 | 1.00 | 1.00 | 1.00 | 1.00 | 1.00 |
| **20** | PL | 1.00 | 1.00 | 1.00 | 1.00 | 1.00 | 1.00 | 0.99 | 1.00 | 1.00 | 0.99 | 1.00 | 1.00 |
| **33** | PL | 1.00 | 1.00 | 1.00 | 1.00 | 1.00 | 1.00 | 1.00 | 0.99 | 1.00 | 1.00 | 1.00 | 1.00 |
| **54** | PL | 1.00 | 1.00 | 1.00 | 1.00 | 1.00 | 1.00 | 1.00 | 1.00 | 0.99 | 1.00 | 1.00 | 1.00 |
| **90** | PL | 1.00 | 1.00 | 1.00 | 1.00 | 1.00 | 1.00 | 1.00 | 1.00 | 1.00 | 1.00 | 1.00 | 1.00 |
| **148** | PL | 1.00 | 1.00 | 1.00 | 1.00 | 1.00 | 1.00 | 1.00 | 1.00 | 1.00 | 1.00 | 1.00 | 1.00 |
| **244** | PL | 1.00 | 1.00 | 1.00 | 1.00 | 1.00 | 1.00 | 1.00 | 1.00 | 1.00 | 1.00 | 1.00 | 1.00 |
| **403** | PL | 1.00 | 1.00 | 1.00 | 1.00 | 1.00 | 1.00 | 1.00 | 1.00 | 1.00 | 1.00 | 1.00 | 1.00 |
| **665** | PL | 1.00 | 1.00 | 1.00 | 1.00 | 1.00 | 1.00 | 1.00 | 1.00 | 1.00 | 1.00 | 1.00 | 1.00 |
| **1096** | PL | 1.00 | 1.00 | 1.00 | 1.00 | 1.00 | 1.00 | 1.00 | 1.00 | 1.00 | 1.00 | 1.00 | 1.00 |
| **1808** | PL | 1.00 | 1.00 | 1.00 | 1.00 | 1.00 | 1.00 | 1.00 | 1.00 | 1.00 | 1.00 | 1.00 | 1.00 |
| **2980** | PL | 1.00 | 1.00 | 1.00 | 1.00 | 1.00 | 1.00 | 1.00 | 1.00 | 1.00 | 1.00 | 1.00 | 1.00 |

### Null models

Table S3: *Mean relative bias of natural ($f$) and general ($\psi$) parameters for the two species null model with weak and strong positive interactions. In the table, we denote >10% bias in bold. To demonstrate how bias changes with sample size, we italise biases between 5-10% and replace values <5% with dots (".").*

| | | High Detection | | | | | | | Low Detection | | | | | | |
|---|---|---|---|---|---|---|---|---|---|---|---|---|---|---|---|
| | | Natural | | | General | | | | Natural | | | General | | | |
| N | Strength | $f_1$ | $f_2$ | $f_{12}$ | $\psi_{00}$ | $\psi_{10}$ | $\psi_{01}$ | $\psi_{11}$ | $f_1$ | $f_2$ | $f_{12}$ | $\psi_{00}$ | $\psi_{10}$ | $\psi_{01}$ | $\psi_{11}$ |
| **20** | Weak | -9.4 | · | **47.12** | -8.12 | · | · | **13.64** | **-102.47** | **-83.69** | **333.51** | **-24.18** | · | -6.68 | **57.22** |
| **33** | | · | · | **40.57** | · | · | · | 7.33 | **-90.3** | **-62.18** | **194.15** | **-20.3** | · | · | **30.98** |
| **54** | | · | · | **17.61** | · | · | · | 5.14 | **-61.66** | **-75.03** | **140.64** | **-18.83** | · | · | **27.96** |
| **90** | | · | · | · | · | · | · | · | **-22** | **-40.65** | **54.16** | **-10.4** | · | · | **14.36** |
| **148** | | · | · | **-11.06** | · | · | · | · | **-15.03** | **-17.33** | **50.08** | -8.53 | · | · | **12.08** |
| **244** | | · | · | -9.92 | · | · | · | · | · | · | **27.93** | · | · | · | 6.52 |
| **403** | | · | · | 7.03 | · | · | · | · | -6.96 | · | 7.25 | · | · | · | · |
| **665** | | · | · | · | · | · | · | · | · | · | **10.97** | · | · | · | · |
| **1096** | | · | · | · | · | · | · | · | · | · | · | · | · | · | · |
| **1808** | | · | · | · | · | · | · | · | · | · | 5.83 | · | · | · | · |
| **2980** | | · | · | · | · | · | · | · | · | · | · | · | · | · | · |
| **20** | Strong | **18.31** | **18.76** | **52.23** | -9 | · | · | 5.23 | **-106.97** | **-99.63** | **83.87** | **-23.91** | **13.58** | 9.83 | 6.73 |
| **33** | | **28.53** | **21.24** | **31.72** | · | · | · | · | · | **-51.63** | **85.31** | **-15.48** | · | 8.69 | 5.5 |
| **54** | | **19.51** | **19.2** | **19.47** | · | · | · | · | **33.58** | **14.04** | **80.11** | **-12.53** | · | 5.12 | 5.41 |
| **90** | | 5.08 | · | 7.15 | · | · | · | · | **61.28** | **38.29** | **78.64** | **-10.54** | · | · | 8.29 |
| **148** | | 5.47 | 8.39 | 5.71 | · | · | · | · | **57.95** | **51.42** | **62.2** | -5.88 | · | · | 6.69 |
| **244** | | 5.66 | · | · | · | · | · | · | **38.32** | **31.2** | **36.77** | · | · | · | 5.64 |
| **403** | | · | · | · | · | · | · | · | 9 | **12.61** | **11.54** | · | · | · | · |
| **665** | | · | · | · | · | · | · | · | 8.61 | 5.08 | 8.73 | · | · | · | · |
| **1096** | | · | · | · | · | · | · | · | · | · | · | · | · | · | · |
| **1808** | | · | · | · | · | · | · | · | · | · | · | · | · | · | · |
| **2980** | | · | · | · | · | · | · | · | · | · | · | · | · | · | · |

### Network complexity

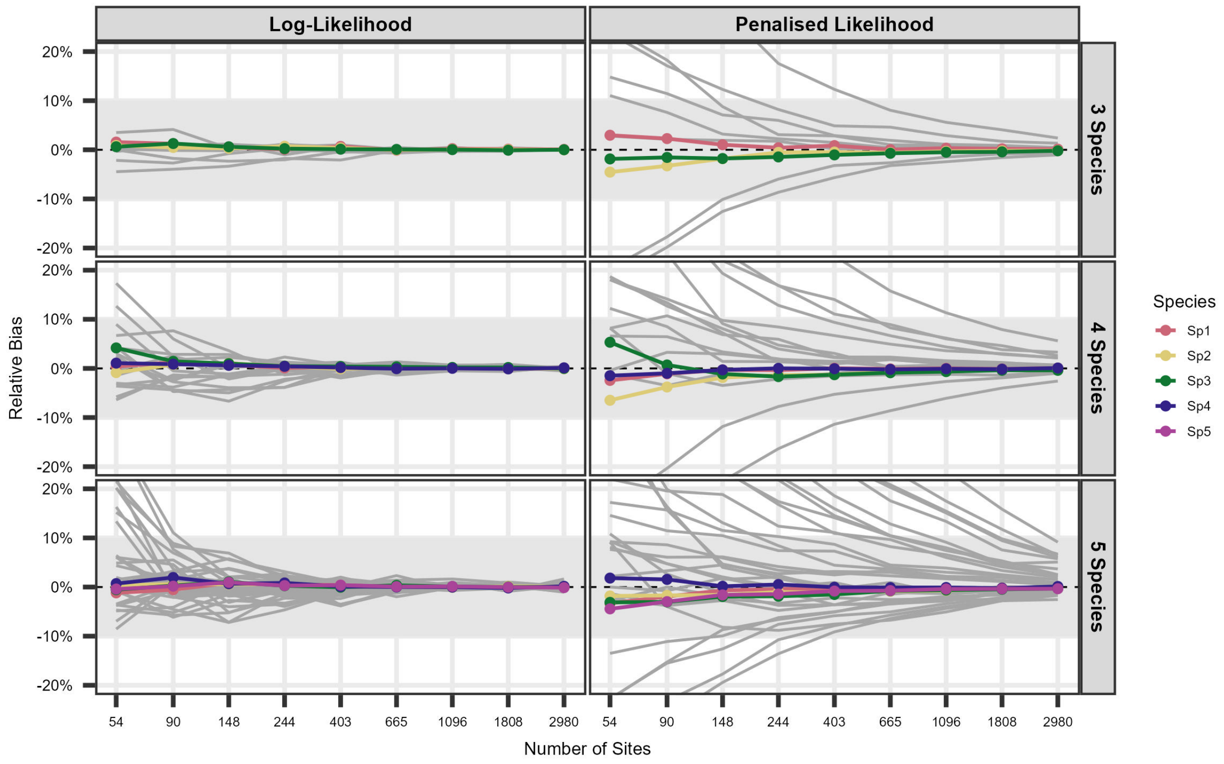


Figure S2: *Relationship between the Relative Bias (%) and sample size for the derived marginal probabilities (colored lines) for all species (1-5) as well as the general parameters (state vector probabilities; grey lines) from the results of the simulated scenarios with increasing network complexity (3-5 species).*

Table S4: *Mean relative bias (%RB) of second-order natural ($f$) parameters in the scenario of 3 and 4 interacting species under normal (LL) and penalized (PL) likelihood estimation methods. In the table, we denote >10% bias in bold. To demonstrate how bias changes with sample size, we italise biases between 5-10% and replace values <5% with dots (".").*

| | | Three species | | | Four species | | | | | |
|---|---|---|---|---|---|---|---|---|---|---|
| N | Likelihood | $f_{12}$ | $f_{13}$ | $f_{23}$ | $f_{12}$ | $f_{13}$ | $f_{14}$ | $f_{23}$ | $f_{24}$ | $f_{34}$ |
| **54** | LL | **32.46** | **15.57** | **30.66** | **38.21** | **33.08** | **24.4** | **22.75** | 9.05 | **16.47** |
| **90** | | **13.52** | 7.89 | **14.84** | **28.08** | · | **17.44** | **23.56** | **15.9** | **16.31** |
| **148** | | 6.23 | · | **10.04** | **16.47** | · | **12.37** | **13.96** | **10.34** | 8.31 |
| **244** | | · | · | · | 6.17 | · | 5.12 | 7.81 | 9.12 | · |
| **403** | | · | · | · | · | · | · | · | · | · |
| **665** | | · | · | · | · | · | · | · | · | · |
| **1096** | | · | · | · | · | · | · | · | · | · |
| **1808** | | · | · | · | · | · | · | · | · | · |
| **2980** | | · | · | · | · | · | · | · | · | · |
| **54** | PL | **-53.15** | **-47.34** | **-60.53** | **-42.45** | **-36.55** | **-46.8** | **-58.29** | **-27.16** | **-46.6** |
| **90** | | **-41.31** | **-32.32** | **-45.2** | **-33.44** | **-30.33** | **-35.99** | **-45.24** | **-15.06** | **-33.74** |
| **148** | | **-23.86** | **-15.49** | **-28.37** | **-20.27** | **-12.72** | **-21.44** | **-32.35** | -5.46 | **-20.53** |
| **244** | | **-14.2** | -6.61 | **-19.84** | **-15.08** | · | **-15.85** | **-22.84** | · | **-15.24** |
| **403** | | -9.01 | · | **-12.4** | **-10.51** | · | **-10.89** | **-17.26** | · | -9.98 |
| **665** | | -5.05 | · | -8.82 | -6.81 | · | -9.29 | **-13.3** | · | -7.04 |
| **1096** | | · | · | -6.27 | -5.3 | · | -6.1 | -9.18 | · | · |
| **1808** | | · | · | · | · | · | · | -7.11 | · | · |
| **2980** | | · | · | · | · | · | · | · | · | · |

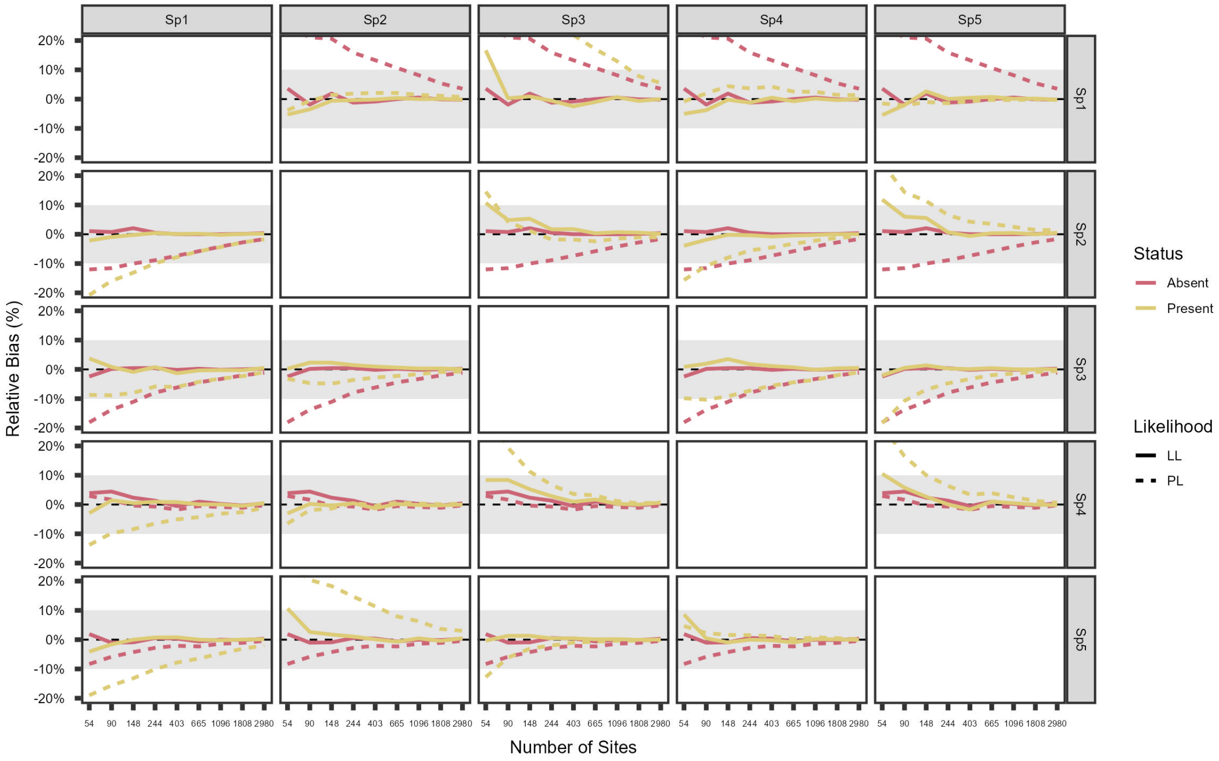


Figure S3: *Relationship between the Relative Bias (%) and sample size for the derived conditional (presence = yellow; absence = red) probabilities from the results of the simulated scenarios with five species (S = 5) in network complexity under log and penalised likelihood.*

11 # Covariate complexity

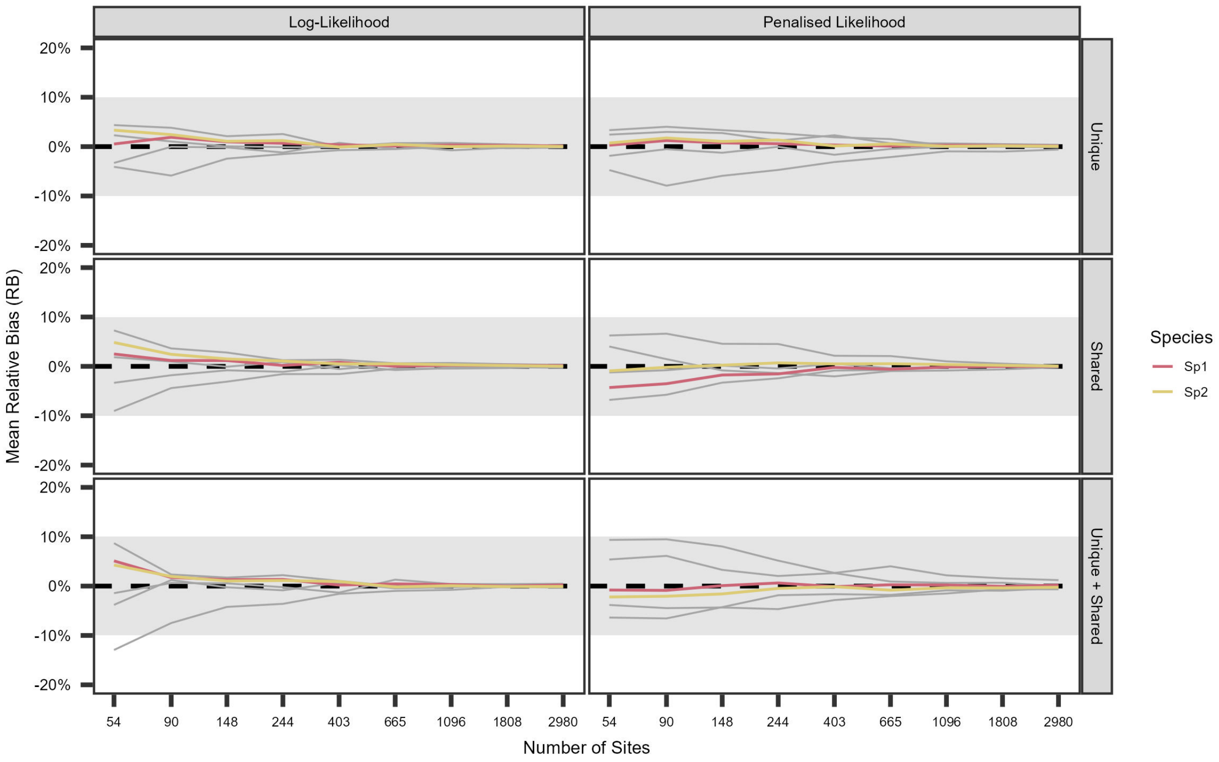


Figure S4: *Relationship between the Relative Bias (%) and sample size (Number of Sites) for the marginal probabilities for Species 1 (red) and 2 (yellow), as well as the general parameters (i.e. state probabilities; grey lines) in scenarios with increasing covariate structure complexity.*

Table S5: *Mean relative bias (%RB) of the conditional probabilities under normal (LL) and penalized (PL) likelihood estimation methods for the covariates complexity scenarios. In the table, we denote >10% bias in bold. To demonstrate how bias changes with sample size, we italise biases between 5-10% and replace values <5% with dots (".").*

<table>
<tr><th></th><th></th><th colspan="4">Unique</th><th colspan="4">Shared</th><th colspan="4">Unique + Shared</th></tr>
<tr><th>N</th><th>Likelihood</th><th>Sp1=1|Sp2=0</th><th>Sp1=1|Sp2=1</th><th>Sp2=1|Sp1=0</th><th>Sp2=1|Sp1=1</th><th>Sp1=1|Sp2=0</th><th>Sp1=1|Sp2=1</th><th>Sp2=1|Sp1=0</th><th>Sp2=1|Sp1=1</th><th>Sp1=1|Sp2=0</th><th>Sp1=1|Sp2=1</th><th>Sp2=1|Sp1=0</th><th>Sp2=1|Sp1=1</th></tr>
<tr><td>**54**</td><td>LL</td><td>.</td><td>.</td><td>.</td><td>.</td><td>.</td><td>.</td><td>.</td><td>.</td><td>.</td><td>.</td><td>.</td><td>.</td></tr>
<tr><td>**54**</td><td>PL</td><td>.</td><td>.</td><td>.</td><td>.</td><td>.</td><td>-6.86</td><td>.</td><td>.</td><td>.</td><td>-6.11</td><td>.</td><td>-7.47</td></tr>
<tr><td>**90**</td><td></td><td>.</td><td>.</td><td>.</td><td>.</td><td>.</td><td>-6.14</td><td>.</td><td>.</td><td>5.35</td><td>-5.79</td><td>.</td><td>-6.74</td></tr>
<tr><td>**148**</td><td></td><td>.</td><td>.</td><td>.</td><td>.</td><td>.</td><td>.</td><td>.</td><td>.</td><td>.</td><td>.</td><td>.</td><td>-5.03</td></tr>
<tr><td>**244**</td><td></td><td>.</td><td>.</td><td>.</td><td>.</td><td>.</td><td>.</td><td>.</td><td>.</td><td>.</td><td>.</td><td>.</td><td>.</td></tr>
</table>

#### Maximum penalty proportion

Fitting with penalized likelihood (PL) is a useful method to circumvent issues with boundary parameter estimates and large biases, common in small sample sizes settings, by shrinking coefficients towards zero by a penalty factor $\lambda$. The optimal value for $\lambda$ is often selected using cross-validation (CV), which can be time-consuming. When fitting the Rota et al. (2016) multispecies occupancy model using the **unmarked** package, the user can use the function *optimizePenalty()* to automate the selection (see Clipp et al. 2021 for a detailed explanation). The default settings propose values from 0.0625 to 16, but using the default range may be inappropriate, requiring stronger (i.e. larger) or weaker (i.e. smaller) penalties respectively. Whilst we used the default settings to speed up the fitting process, we highlight below the frequency of encountering penalties at the minimum-maximum default settings when increasing the number of species or introducing covariates.

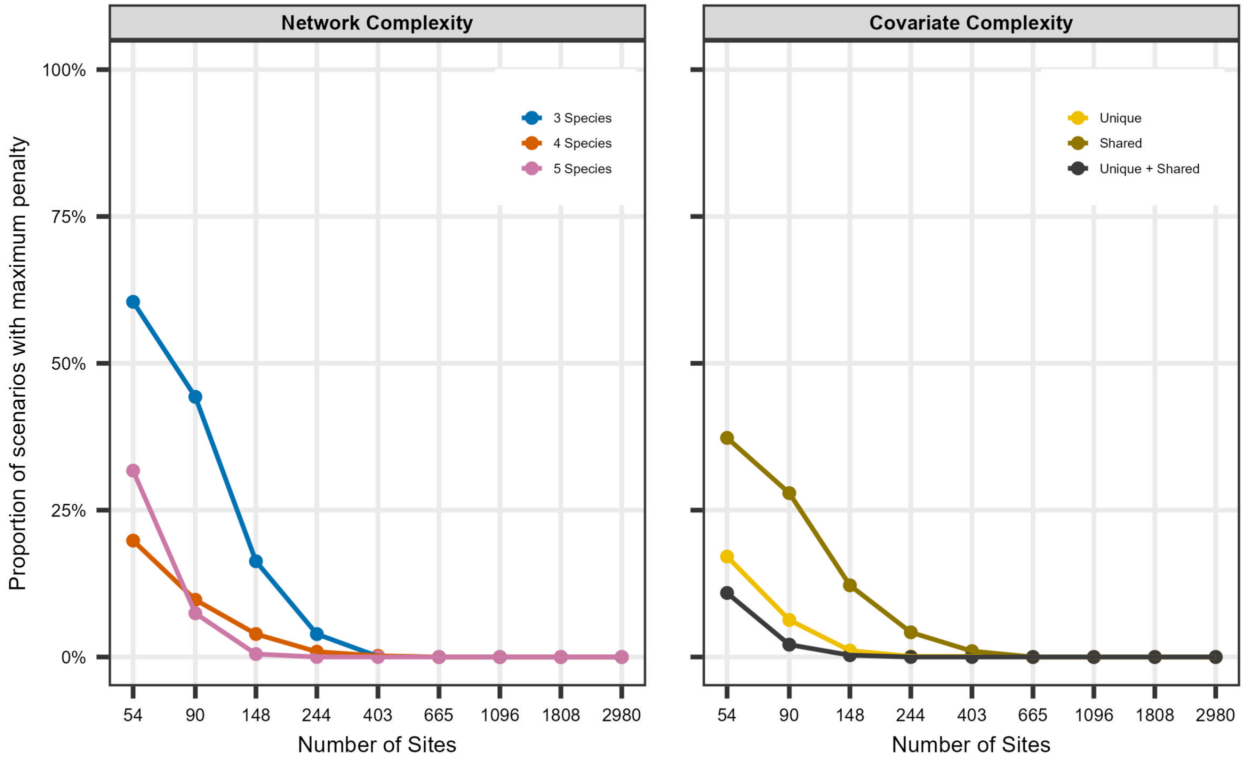


Figure S5: *Relationship between the proportion (%) of network and covariate complexity scenarios with minimum ($\hat{\lambda} = 0.0625$) or maximum ($\hat{\lambda} = 16$) default values for the penalty term when fitting the simulated models using penalized likelihood.*

The occurrence of extreme penalties declines sharply as sample size increases, with no issues observed for sample sizes above 400 (Figure S5). Increasing the number of species (Network) tends to produce extreme penalties more frequently than increasing model complexity via covariates (Covariate). When using default settings, users can expect extreme penalties to occur 10–60% of the time for sample sizes below 100. It is recommended that users examine the $\hat{\lambda}$ estimates and adjust the values as needed, especially if they are working with fewer than 100 sites. Without appropriate consideration of the selection penalty values, users might be unknowingly introducing more bias in cases where the extreme default values are used for evaluations of co-occurrence patterns (Figure S6).

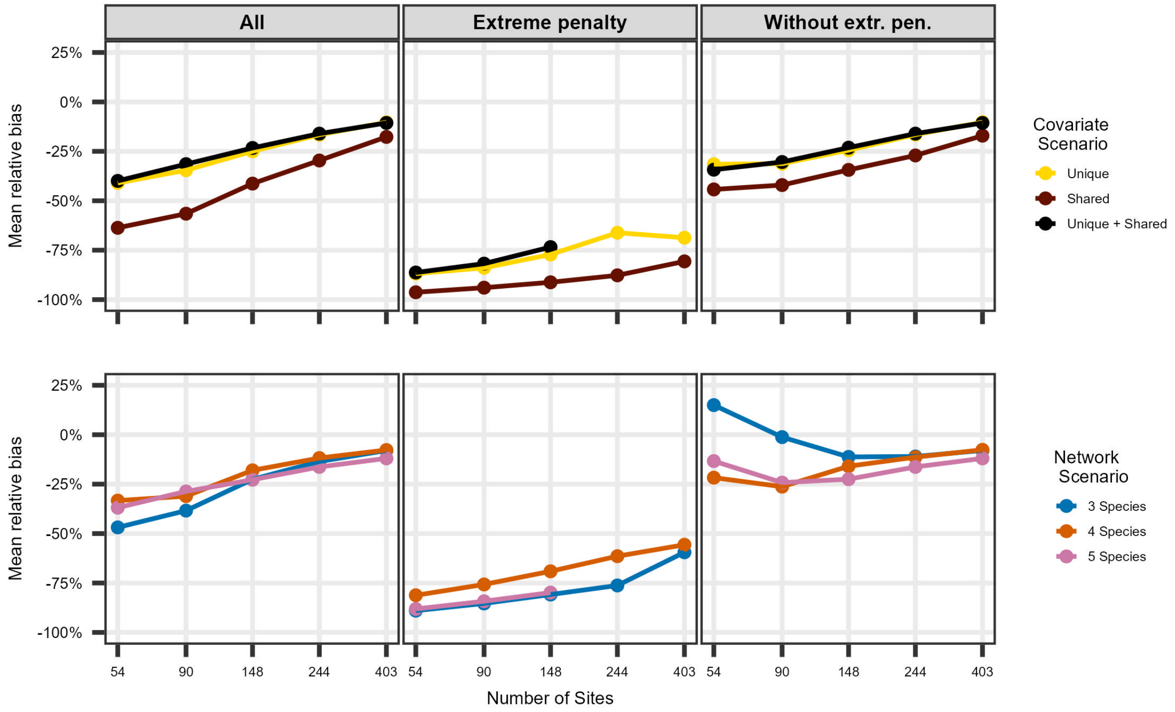


Figure S6: *Comparison of the relationship between the mean relative bias (%) and sample size across all second order natural parameters for each network and covariate complexity scenarios fitted under penalized likelihood usng: all simulations (left); only those minimum ($\hat{\lambda} = 0.0625$) or maximum ($\hat{\lambda} = 16$; i.e. extreme) penalties; and scenarios without the extreme (extr.) penalty values (right). Relative bias was larger across all scenarios and sample sizes irrespective of scenario, highlighting the need of evaluation when using penalized likelihood methods using the default settings. Sample sizes above 403 were not evaluated as they had no extreme penalty values (Figure S5)*
